# Constructing early diagnosis model of colorectal cancer based on expression profile

**DOI:** 10.1101/2020.01.06.895730

**Authors:** shaoqiang Wang, Yifan Wang, Shudong Wang

## Abstract

**purpose:** In order to break through the restrictive factors such as the fecal occult blood test (FOBT) in the routine detection of bowel cancer, which is susceptible to diet and drugs, and the high cost and inconvenience of microscopy, Seeking a possible FOBT alternative.

**Methods:** An error back propagation neural network (BPNN) algorithm was used to construct a CRC diagnosis model based on expression profiles.

**Results:** The accuracy of the model on the training and test sets is 0.943 and 0.935, respectively. AUC all reached above 0.95.

**Conclusion:** The CRC molecular detection model based on expression profiles provides a possible alternative to FOBT. It provides a new approach and method for the clinical diagnosis of bowel cancer.

## 0 background

Colorectal cancer (CRC) is a collective term for colon and rectal adenocarcinoma. CRC has a high incidence worldwide, and its mortality rate is also among the leading causes of tumor death.Compared with other tumors, CRC has higher heterogeneity, so it can be divided into subtypes with different characteristics based on clinical or molecular characteristics. The cause of CRC is more complicated, but whether it is primary CRC (~ 70%) or hereditary CRC (10-25%), it shows specific characteristics at the molecular level [4–6], including the genome Chromosomal instability (CIN), loss of heterozygosity, and copy number variation. Epigenetic changes, such as CpG island methylation [26], have been shown to drive adenomas to cancer in a sporadic and genetic form of CRC.Because CRC develops slowly in precancerous lesions, it is usually in the middle and late stages of CRC when patients are aware of it, and this has a great impact on the prognosis of CRC, so early detection can reduce the incidence and mortality of CRC.Considering the high diagnostic performance, optical colonoscopy (OC) is the gold standard study for early detection of CRC. Colonoscopy can be performed concurrently with biopsy specimens for a clear diagnosis, and at the same time as a therapeutic polypectomy, thus preventing long-term CRC death [1]. However, patients with tumor-related stenosis, older patients and those with comorbidities are more likely to have incomplete or difficult optical colonoscopy [12,13]. Pickhardt et al. [7]found that CTC was comparable to colonoscopy in detecting larger colon polyps.Two meta-analytical studies have shown that carbon tetrachloride has a 87.9 % high sensitivity (100%) for detecting colon cancer and is less than 10 mm for adenomas [8,9].Despite such encouraging data, there is currently no cross-continental consensus on whether CTC should be used as a screening method for asymptomatic patients. Detection based on genomic mutations [10–12]Although it provides great help for accurate diagnosis and targeted therapy of CRC, the high heterogeneity of CRC limits the use of this method. MRI [15]is the recommended method for initial staging, because its definition of localization determines the overall expansion of the tumor and its relationship with the peritoneal reflex with high accuracy. Tumor growth on the initial MRI should be best described in relation to anatomical structures such as the mesorectal fascia [16]. Most of the staging failures of MRI occur in the differentiation of T2 and marginal T3 stages, and staging is the main cause of error [17]. Although previous studies have not shown the great advantages of dedicated phased array coils [18], our clinical experience is positive, and in our institution we use phased array coils as the standard for the initial diagnosis of colorectal cancer. The advantage of high spatial resolution with a large field of view is that phased array MRI is suitable for staging superficial and advanced rectal tumors. Endorectal ultrasonography (ERUS) is now the established model for assessing rectal wall integrity.The accuracy of T staging is between 69% and 97%.Intrarectal ultrasound (US) is currently the most accurate imaging method for assessing T1 tumors [18]. ERUS and intrarectal MRI have similar accuracy in distinguishing superficial (T1 and T2) and T3 tumors [20]. However, intrarectal MRI is associated with high costs, limited availability, and patient discomfort. Therefore, the European Medical Association guidelines do not recommend intrarectal MRI as the preferred imaging method for clinical T stage of colorectal cancer [15].

Conventionally, methods such as fecal occult blood test (FOBT) [13,14]and colonoscopy are also used to diagnose CRC. Although both have certain advantages, FOBT is also vulnerable to diet and drugs, and Limiting factors such as the high cost and inconvenience of microscopy. In this study, an error back propagation neural network (BPNN) algorithm [27–31]was used to construct a CRC diagnosis model based on expression profiles. The relationship between gene expression changes and CRC and the possibility of CRC diagnosis were explored. CRC molecular detection of expression profiles [21–25]provides a possible alternative to FOBT.

## 1 Analytical method

### 1.1 Data source and preprocessing

The colorectal cancer (CRC) dataset we used was derived from TCGA (The Cancer Genome Altas) and GEO (Gene Expression Omnibus). First use the GDC Data Transfer Tool to download the RNA seq data (read count) of colon adenocarcinoma (COAD) and rectal adenocarcinoma (READ) from the TCGA (https://portal.gdc.cancer.gov/) database and the clinical data corresponding to the sample information. According to the COAD and READ information recorded by the TCGA, 41 pairs of COAD normal and tumor samples and 10 pairs of READ normal and tumor samples were obtained. Then download the CRC gene expression profile data from the GEO database (https://www.ncbi.nlm.nih.gov/geo/), including GSE39582 (19 normal samples, 443 tumor samples), GSE41258 (54 normal samples), 186 tumor samples) and GSE44076 (98 normal samples, 50 mucosa samples, 98 tumor samples). In order to ensure the consistency of the differential expression analysis of different data sets, we downloaded the raw data of GSE39582, GSE41258, and GSE44076, respectively, and used the RMA (robust mean analysis) method for homogenization. The pre-processed sample information is shown in Table 1, and finally contains 1049 CRC samples for subsequent analysis.

**Table 1.** Sample information statistics

|  | GSE39582 | GSE41258 | GSE44076 | TCGA-COAD | TCGA-READ |
| --- | --- | --- | --- | --- | --- |
| Platform | hgu133plus2 | hgu133a | hgu219 | NGS | NGS |
| Tumor | 443 | 186 | 98 | 41 | 9 |
| Mucosa | 0 | 0 | 50 | 0 | 0 |
| Normal | 19 | 54 | 98 | 41 | 10 |
| DataType | tumor+adjacent | tumor+adjacent | cancer+healthy | tumor+adjacent | tumor+adjacent |

### 1.2 Differentially Expressed Gene Screening

Differentially expressed gene analysis is mainly based on GSE39582 and GSE41258 data, because both are affymetrix platforms and the sample type is tissue. The limma (version 3.8) tool was used to identify differentially expressed genes (DEGs) in normalized and tumor samples of GSE39582 and GSE41258 after homogenization. Genes with fold change more than 2 times and FDR (BH adjusted P-value) <0.05 were taken as DEGs. TCGA’s COAD and READ data are of the NGS type. The read count value of the transcript was analyzed by DESeq2, and the FDR <0.05 DEGs threshold was also taken. Before the above differential expression analysis, the similarity of the samples was evaluated on the GEO and TCGA datasets, and the correlation coefficients between the samples were calculated. The results show that the tumor and normal samples from different sources have high internal consistency (**S1_Fig**). Both the heatmap and volcano map expressed by DEGs were constructed using R software.

### 1.3 Functional enrichment analysis

We use clusterProfiler (version 3.8) package to perform DEA on the biological process (BP) of Gene Ontology (GO), cellular component (CC), molecular function (MF) and KEGG pathway enrichment analysis. Take q value <0.05 as the threshold for significant enrichment. The dotplot of clusterProfiler displays the enrichment result.

### 1.4 protein interaction analysis

Using the protein interaction (PPI) information provided by the STRING database ((https://string-db.org/), we built a PPI network of DEGs, retaining PPI information with confidence socre> 0.9, and using Cytoscape (version 3.7.1) Show PPI network. Hub gene analysis uses Cytoscape’s NetworkAnalyzer plug-in for analysis, calculates the connectivity degree of each gene (node), and ranks genes according to the connectivity degree. For genes that are significantly up-regulated and significantly down-regulated, construct the above network and take the degree respectively The largest gene is determined as the hub gene, and finally 2 hub genes are obtained. The PPI network module analysis uses the MCODE tool and the parameters take the default value. The GO and KEGG analysis of the genes in the module also use the clusterProfiler tool.

### 1.5 Construction of a neural network-based diagnostic model

Using the error back propagation neural network (BPNN) algorithm, we constructed a CRC diagnosis model based on hub genes. First, randomly group the Normal (healthy), Mucosa, and CRC samples of the GSE44076 dataset, set seed = 12345, and divide the 246 samples into a training set and a testing set evenly. The main parameters of the BPNN algorithm are the learning rate, the lambda of the regular term coefficient, the number of hidden layers, and the number of neurons included in the hidden layer. In order to find the optimal parameters, a grid search method is used to evaluate the performance of the model under different parameter combinations. Since our target value is a categorical variable, the accuracy of the model prediction is used as the model’s judgment index. Accuracy is calculated as follows:

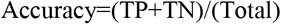

Finally, the model with the maximum training set and testing set accuracy (training set accuracy + testing set accuracy-1) is the optimal model. The model parameter learning rate = 0.006, lambda = 6e-04, hidden layer = 10 neurons. To avoid biasing the model by random grouping, we used the bootstrap method to calculate the accuracy (S12_Table) of the training set and testing set of the model under 100 random samples.

### 1.6 Statistical Analysis

Statistical analyses were performed using R (version 3.5.2) software. Student t-test was used to test the significance of differences in gene expression levels of paired samples, and Wilcox rank test was used to perform a two-group significance test of gene expression levels of unpaired samples. The Kruskal-Wallis rank test was used for the significance test of two or more groups, and the FDR was calculated using the BH-method. In this study, unless otherwise specified, *** indicates p <1e-5, ** indicates p <0.01, and * indicates p <0.05.

## 2 Numerical Simulation

### 2.1 Analysis process

In this study, the gene expression profile data of normal and tumor samples provided by the GSE39582 and GSE41258 datasets were first used to calculate the differentially expressed genes (DEGs) of the two using the limma tool. The DEGs common to both were used for subsequent verification. Using the TCGA’s COAD and READ data sets, we verified the identified DEGs to further determine the reliability of our DEGs. We then commented on the possible functions of DEGs, including participating biology processes (BP) and pathways. Analysis of protein interactions allows us to have a deeper understanding of changes in cellular pathways (signaling pathways, metabolic pathways) that may be involved in the transition from normal to tumor.

### 2.2 (DGEs) Identification of differentially expressed genes

Using the limma tool, we analyzed the differentially expressed genes in the normal and tumor grouped samples in the GSE39582 and GSE41258 datasets, and obtained 1691 (up / down: 689/1002) and 414 (up / down: 111/303) differentially expressed genes (Fig 1A-B, S1-S2_Table). After removing genes with inconsistent expression patterns in the two sets of data sets, a total of 270 DEGs (up / down: 90/180, Fig 1C, S3_Table) were obtained. The total DEGs accounted for 19.6% and 76.3% of the total DEGs, respectively. The large number of DEGs indicates that the occurrence of CRC involves more molecular-level changes. Further analysis of 270 common DEGs revealed that 90 genes were significantly up-regulated in the tumor group and 270 genes were significantly down-regulated in the tumor group (Fig 1D-E), indicating that the activation and inhibition of certain biological processes may be involved The CRC normal transitions to the tumor state.

**Fig 1.**
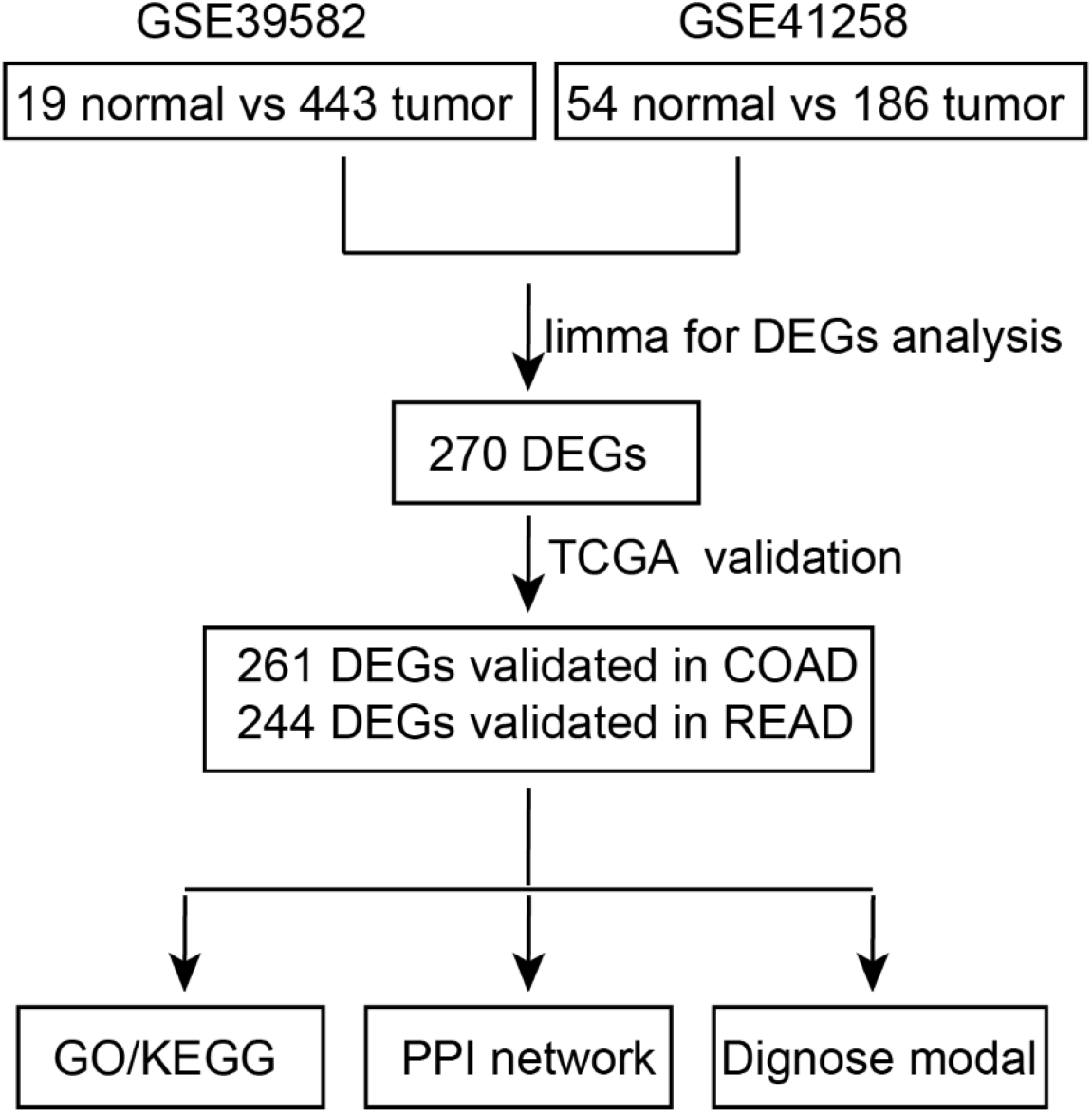
Research flow chart. DEGs: different expression genes, COAD: colon adenocarcinoma, READ: rectal adenocarcinoma, PPI: protein-protein interaction_o_

### 2.3 DEGs Functional Analysis

In order to further study the functions of these DEGs, we performed GO function annotation on 270 DEGs. Due to the significant difference in up- and down-regulated gene expression patterns, we analyzed the function of up- and down-regulated genes, respectively. We see that genes that are up-regulated in the tumor are mainly involved in biological processes related to the extracellular matrix and extracellular structure (Fig 2A, S4_Table), while the genes that are down-regulated are involved in biological processes that are significantly different from up-regulated genes, mainly related to ion Detoxification is related to stress response (Fig 2B, S5_Table). Analysis of the KEGG pathway found that genes that were significantly up-regulated in the tumor were significantly enriched in the ECM-receptor interaction, focal adhesion, and PI3K-Akt signaling pathway (Fig 2C, S6_Table), and these pathways are importantly related to tumor formation and progression. The down-regulated genes were significantly enriched in the pathways such as Fatty acid degradation, Glycolysis / Gluconeogenesis (Fig 2D, S7_Table), which indicates that these genes involved in fatty acid and glucose metabolism are inhibited in tumor cells.

**Fig 2.**
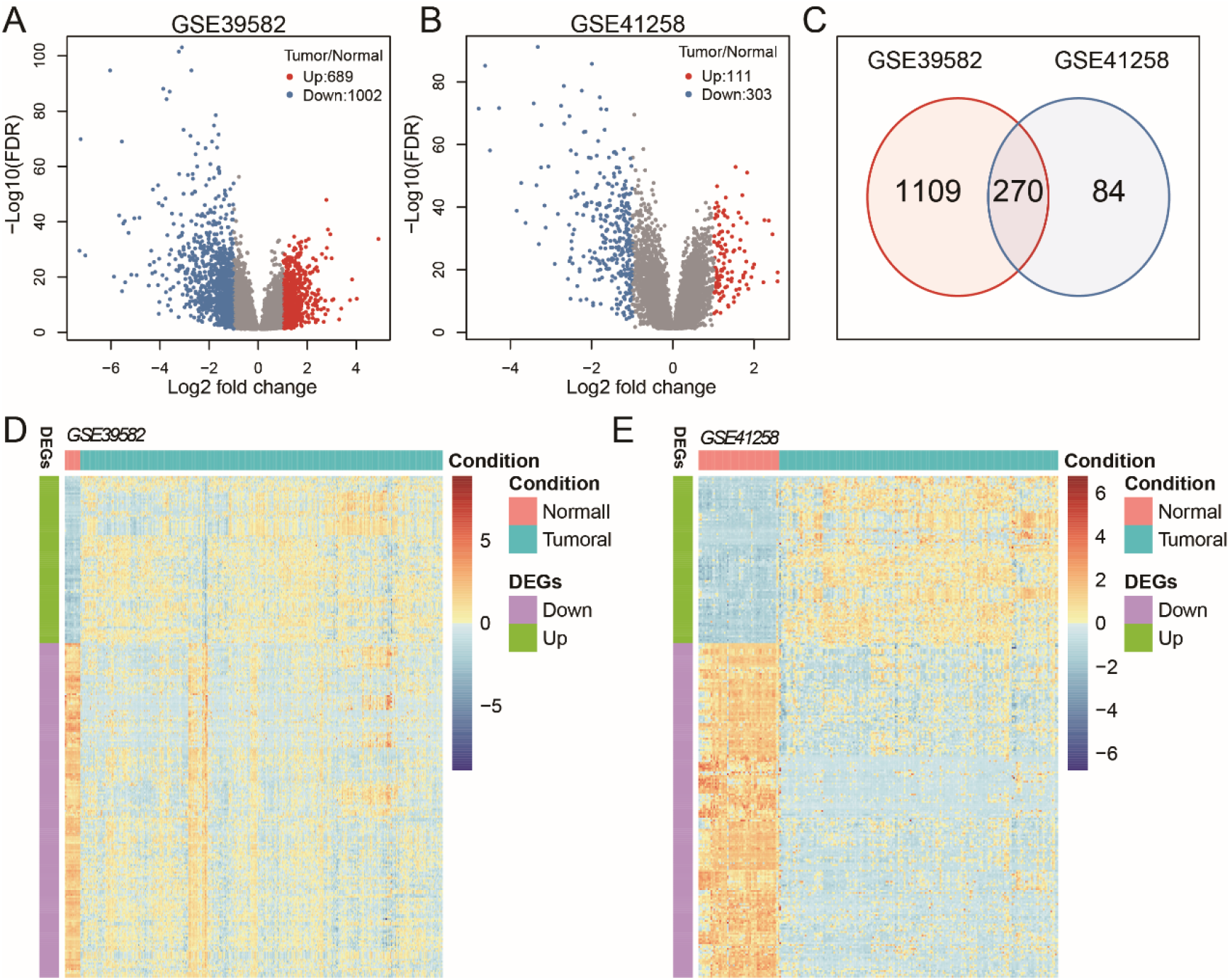
Identification of differentially expressed genes A: GSE39582 Fold change of differentially expressed genes in the dataset and volcano plots of FDR; B: Volcano plot of fold change and FDR of differentially expressed genes in the GSE41258 dataset; C: GSE39582 and GSE41258 datasets differential expression overlap; D: Heat map of shared differentially expressed genes on the GSE39582 dataset; E: The shared differentially expressed genes are heat map expressed on the GSE41258 dataset.

### 2.4 DEGs Interaction Network and Hub Gene Analysis

The 90 DEGs up and 180 DEGs down get 715 and 1089 PPI network edges, respectively. These edges have a confidence score> 0.9. Hub gene analysis found that the CCND1 and FOS genes had the highest degree of up-regulation and down-regulation in the DEGs network and were significantly higher than other genes (43/54, S8_Table, S9_Table). The module analysis can significantly divide the up-regulated DEGs network into 3 subclusters, which contain 25, 12, and 5 genes, respectively. Among them, subcluster1 is closely related to cancer occurrence, and the functions of subcluster2 and subcluster3 are unknown (Fig 4A, S8_Table). The down-regulated DEGs network can be significantly divided into 5 sub-networks, among which subcluster1 is related to glycolysis / gluconeogenesis metabolism, subcluster2 is related to ion metabolism, subcluster3 is related to bile secretion, and subcluster4 is related to nitric biosynthesis process (Fig 4B, S9_Table).

**Fig 3.**
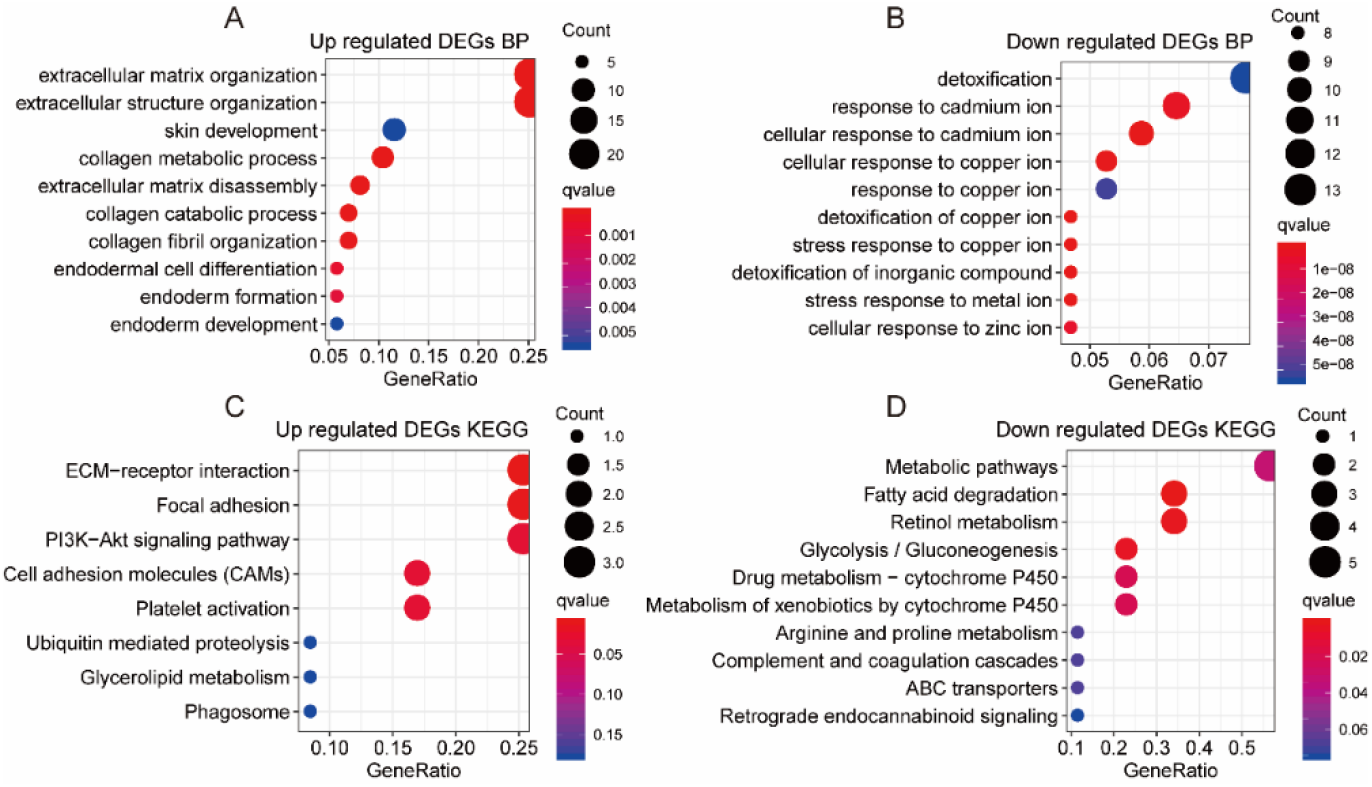
GO and KEGG annotation of differentially expressed genes (show top 10 categories). A: GO enrichment results for shared up-regulated genes; B: GO enrichment results for shared down-regulated genes; C: KEGG pathway enrichment results for shared up-regulated genes; D: KEGG pathway enrichment results for the shared down-regulated genes.

**Fig 4.**
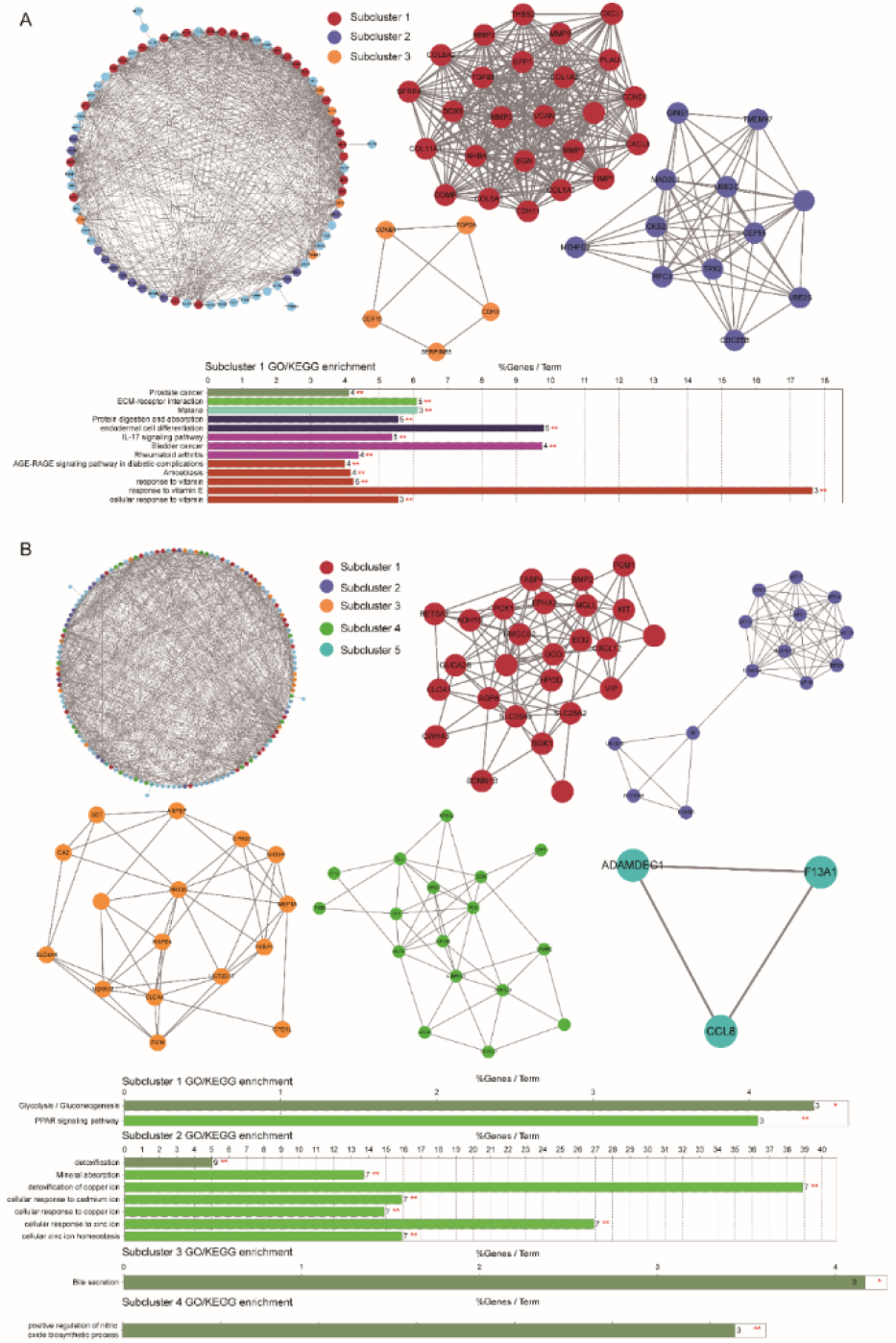
Analysis of Differentially Expressed Gene Interaction Networks A: Module Analysis and Function Notes for Upgrading DEGs Network; B: Modified the module analysis and function notes of DEGs network.

### 2.2 DEGs expression analysis in independent validation set

Using expression data of intestinal cancer (colon and rectal adenocarcinoma) provided by TCGA, we verified the expression of 270 DEGs. Since TCGA’s CRC expression data is of RNA seq type, we used the read count values corresponding to these DEGs to compare the overall expression of up- and down-regulated genes on normal and tumor. It is significantly higher than normal, and the down-regulated genes are also significantly lower than normal on the tumor (Fig 5A-B). These are highly consistent with our results based on the GEO chip expression data. Further testing the expression levels of each of the up-and down-regulated genes in tumor and normal, it was found that in COAD and READ samples, 99% (261/263) and 93% (244/263) of the gene expression were significant. The difference (FDR <0.05, Fig 5C-D, S10-S11_Table), which further shows the reliability of the DEGs we identified.

**Fig 5.**
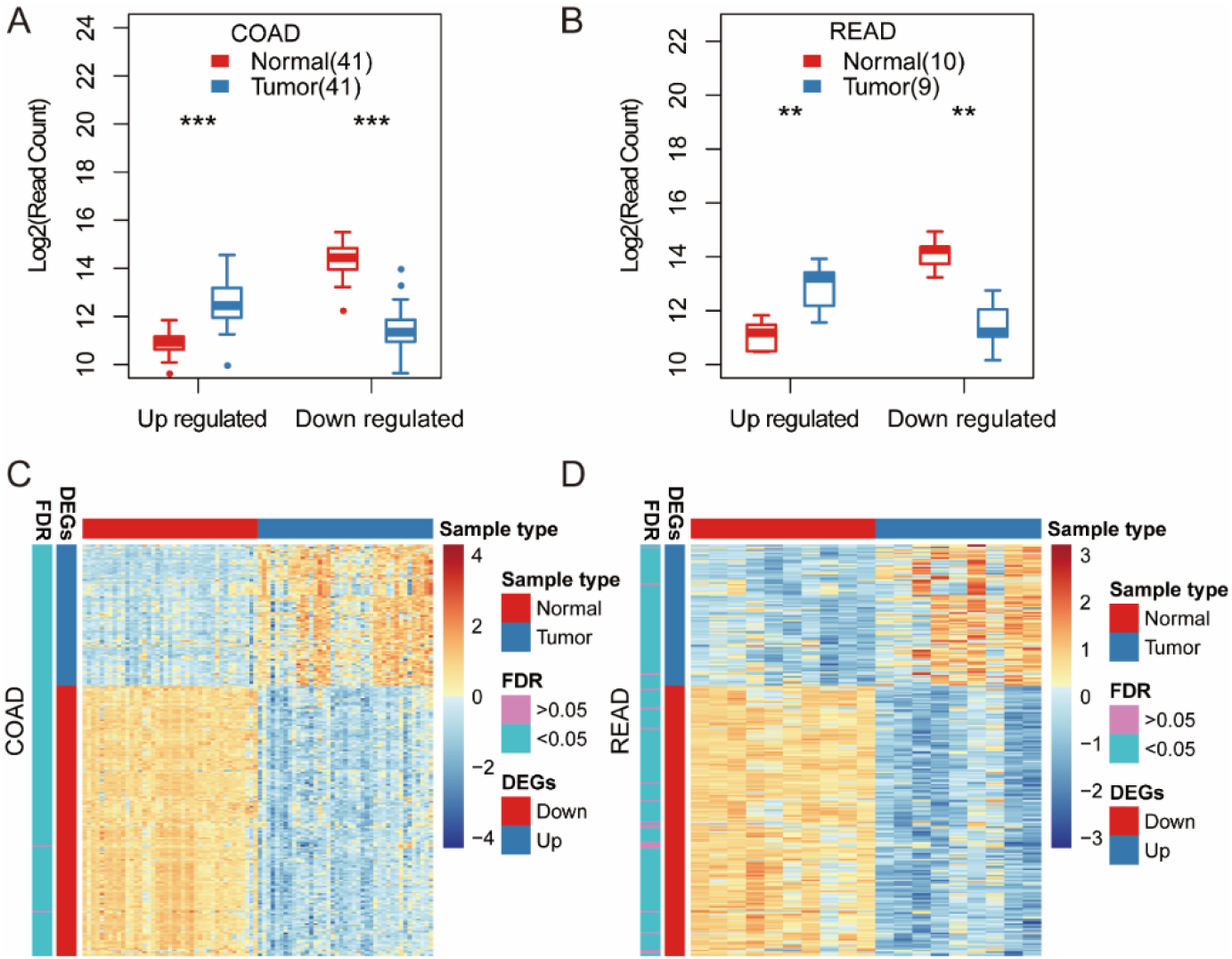
The expression of differentially expressed genes on the TCGA CRC dataset. A: Up- and down-regulation of overall gene expression levels in tumor and normal samples of COAD; B: Up- and down-regulation of overall gene expression levels in READ’s tumor and normal samples; C: Differential expression test of up- and down-regulated genes in tumor and normal samples of COAD; D: Up-regulated and down-regulated genes were tested for differential expression in tumor and normal samples of READ.

### 2.6 Constructing a neural network-based diagnostic model

The neural network model constructed based on the expression values of FOS and CCND1 of the two hub genes has an accuracy of> 0.9 on both the training set and the testing set, and the median accuracy of 100 random samples reached 0.943 and 0.927, respectively (Fig 6A, S11_Table), indicating random grouping It has less impact on our model. On the whole, the AUC predicted by the model for Normal (healthy), Mucosa, and CRC samples also exceeded 0.97 (Fig 6B-C), indicating that the prediction model based on the FOS and CCND1 genes has good performance. Furthermore, we compared the expression levels of the two genes in all samples, and there was no strong correlation between the expression levels of the two genes (cor = 0.16, p = 0.013). The expression levels of these two genes in Mucosa samples were significantly lower than those in Normal (healthy) and CRC samples (p <1e-5), while the FOS gene expression in Normal (healthy) samples was significantly higher than that in CRC samples, and the expression characteristics of CCND1 gene were opposite. It is significantly higher than Normal on CRC (Fig 6D).

**Fig 6.**
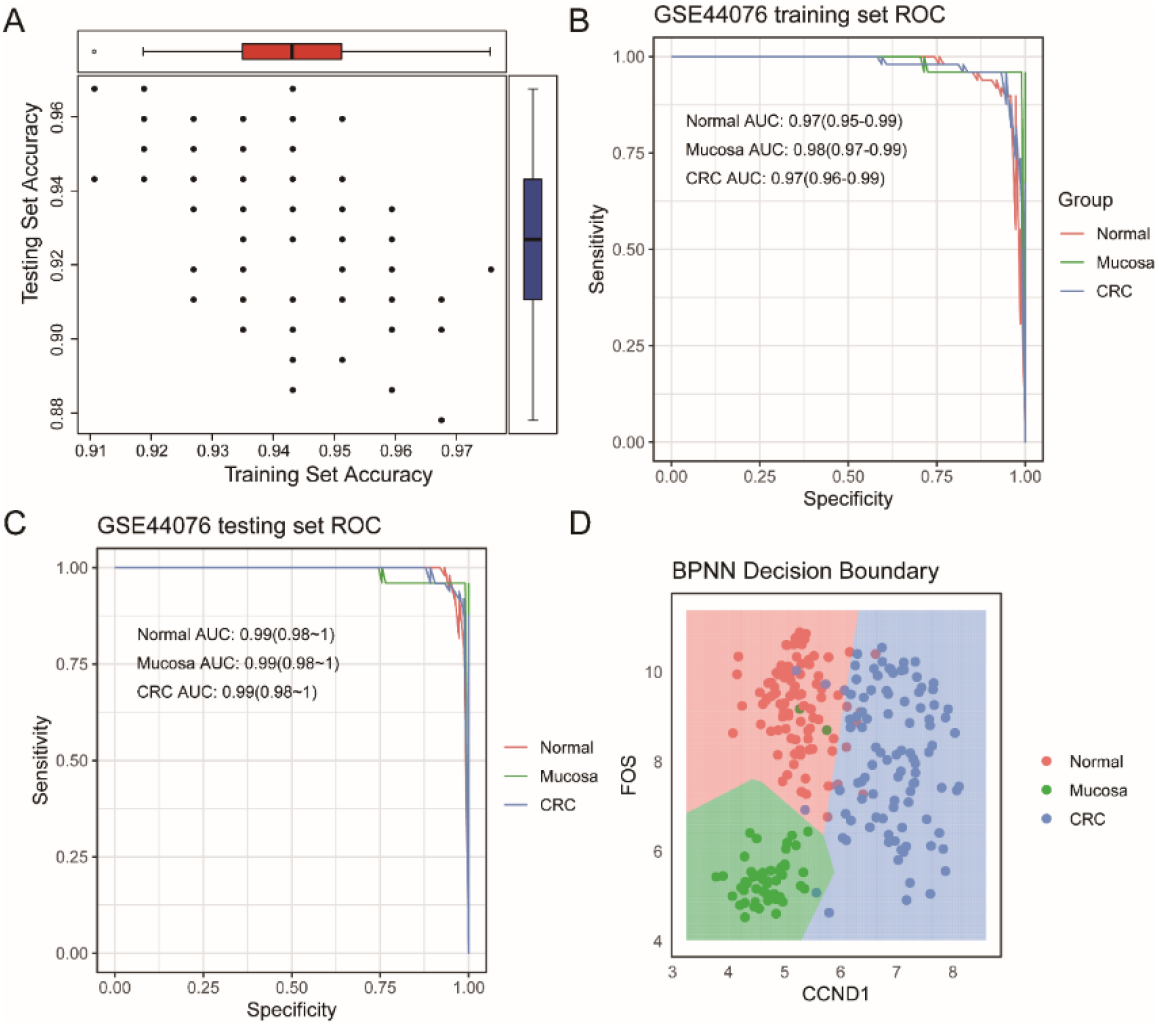
CRC diagnostic model based on FOS and CCND1 genes. A: 100 times random training and test set accuracy distributions; B: Model ROC and AUC values of training set; C: Model ROC and AUC values of testing set; D: FOS and CCND1 genes were distributed on Normal (healthy), Mucosa and CRC samples.

## 3 In conclusion

1. This study used the GSE39582 and GSE41258 data sets from GEO to identify a group of differentially expressed genes between normal and CRC. A total of 270 DEGs were obtained on two sets of data sets from different sources, 90 of which were in CRC samples. Medium up-regulated genes and 180 down-regulated genes in CRC samples.
2. TCGA database’s CRC (colon adenocarcinoma + rectal adenocarcinoma) independent data set was used to verify 270 DEGs. Among them, more than 90% of the genes showed differential expression in normal and tumor samples, and the expression patterns were up- and down-regulated Also consistent. This shows that our DEGs filtered based on the GEO dataset are reliable.
3. The functional annotation of differentially expressed genes found that genes that are up- regulated in the tumor are mainly involved in biological processes related to the extracellular matrix and extracellular structure, while down-regulated genes are mainly related to the detoxification and stress response of ion. Pathway enrichment analysis shows that the pathways involved in upregulated genes are mainly related to tumor formation and development, while the pathways involved in downregulated genes are mainly related to fatty acid and sugar metabolism.
4. Analysis of the protein interaction network based on DEGs shows that the hub genes with a degree significantly higher than other genes: FOS and CCND1, where the FOS gene is the hub gene that down-regulates the DEGs network, and CCND1 is the hub gene that up-regulates the DEGs network. The module analysis of the interaction network divides the up- and down-regulated DEGs networks into 3 and 5 sub-networks, respectively. The functions of the subclusters are significantly different.
5. Using two hub genes: FOS and CCND1, we constructed a CRC diagnostic model based on the neural network algorithm. The accuracy of the model on the training set and the test set was 0.943 and 0.935, respectively, and the AUC reached above 0.95, reflecting our The model has better performance.

## ethics approval and consent to participate

This article does not contain any studies with human participants or animals performed by any of the authors.

## Consent for publication

Not Applicable

## The individual contributions of authors

WYF collected and analyzed the feasibility of the data and revised the paper.WSQ uses software to analyze the data and compare the results, and was a major contributor in writing the manuscript.WSD checks the direction of the paper and provides guidance on the selection of methods and the establishment of models.All authors read and approved the final manuscript.

## Acknowledgements

Not Applicable.

## Abbreviations

CRC: The colorectal cancer;
TCGA: The Cancer Genome Altas;
GEO: Gene Expression Omnibus;
FOBT: fecal occult blood test;
BPNN: error back propagation neural network;
COAD: colon adenocarcinoma;
READ: rectal adenocarcinoma;
DEGs: differentially expressed genes;
BP: biological process;
GO: Gene Ontology;
CC: cellular component;
MF: molecular function;

## The Availability of data and materials

The colorectal cancer (CRC) dataset we used was derived from TCGA (The Cancer Genome Altas) and GEO (Gene Expression Omnibus). The datasets used and/or analysed during the current study available from the corresponding author on reasonable request.

---

**S1 Fig.**
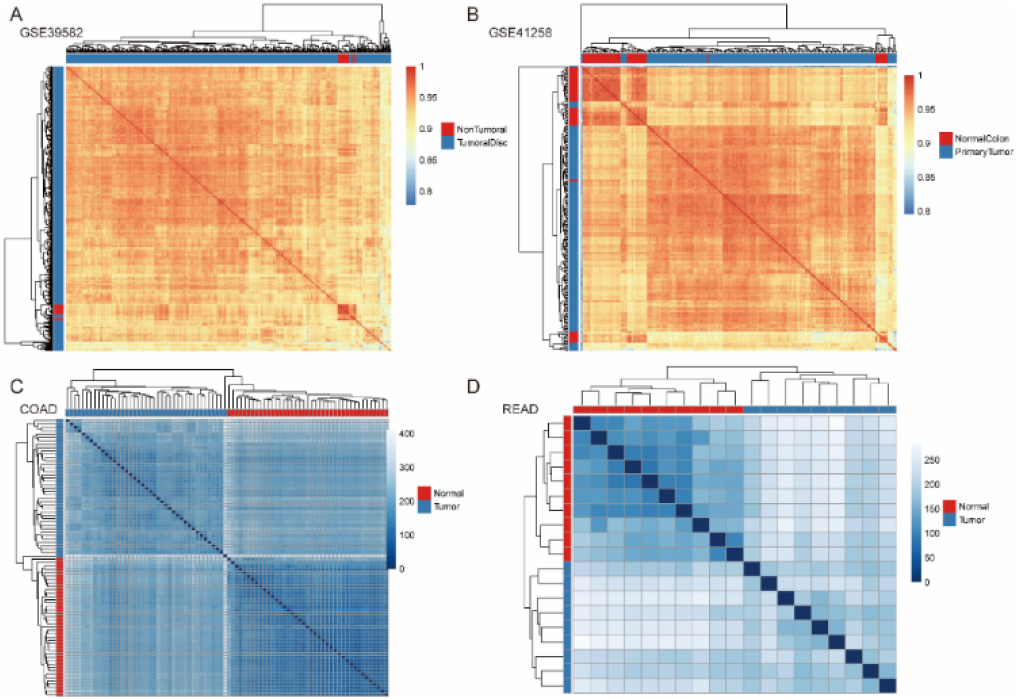
Similarity evaluation of Tumor and normal samples.

